# A unifying model of LAT condensates in reconstitution experiments

**DOI:** 10.1101/2025.10.31.685772

**Authors:** Yannick A. D. Omar, Simou Sun, Mehran Kardar, Jay T. Groves, Arup K. Chakraborty

**Affiliations:** Department of Chemical Engineering, Massachusetts Institute of Technology, Cambridge, MA 02139, USA; Department of Chemistry, Stony Brook University, Stony Brook, NY, USA; Department of Physics, Massachusetts Institute of Technology, Cambridge, MA 02139, USA; Department of Chemistry, University of California, Berkeley, CA, United States; California Institute for Quantitative Biosciences, University of California, Berkeley, CA, United States; Institute for Medical Engineering and Science, Massachusetts Institute of Technology, Cambridge, MA 02139, USA; Ragon Institute of Massachusetts General Hospital, Massachusetts Institute of Technology and Harvard University, Cambridge, MA 02139, USA; Department of Chemistry, Massachusetts Institute of Technology, Cambridge, MA 02139, USA

## Abstract

The formation of condensates by the Linker for the Activation of T-cells (LAT) is a key signal gating and amplification step in the T-cell receptor signaling pathway. LAT condensation is challenging to study in-vivo and is therefore often investigated using reconstitution experiments. While these experiments recapitulate key aspects of LAT condensation, they also exhibit some puzzling features. Here, we describe the mechanisms underlying these observations using two complementary models. First, we employ a Smoluchowski aggregation model to show that the delay time before condensation is observed arises from a low effective binding probability between LAT monomers. Second, we propose a field-theoretic model that reproduces all condensate morphologies observed in experiments, showing that they can arise from common underlying dynamics modulated by variations in experimental conditions. This result unifies different experimental observations reported previously. While this article addresses open questions regarding the formation of LAT condensates, our results also provide a common framework for understanding condensation of other multivalent membrane proteins such as EGFR, FGFR2, and nephrin.

## Introduction

T-cells display T-cell receptors (TCRs) on their surface that recognize complexes of pathogen-derived peptide-major histocompatibility (pMHC) molecules on antigen-presenting cells (APCs) with high specificity and sensitivity. Identification of foreign pMHC can then lead to T cell activation, with activated T cells playing a critical role in mounting an effective cell-mediated immune response. Robust recognition of foreign pMHC is achieved through a sequence of biochemical, non-equilibrium kinetic proofreading steps (***McKeithan, 1995***; ***Ganti et al., 2020***) that are followed by the adapter protein LAT (Linker for Activation of T cells) forming a condensate. This condensate functions as a scaffold for key signaling complexes (***Wange, 2000***) and thus facilitates relaying the recognition of foreign pMHC to downstream pathways such as NFAT and NF-*κ*aB translocation into the nucleus (***Hogan et al., 2003***; ***Thaker and Rudd, 2015***; ***McAffee et al., 2022***; ***Morita et al., 2024***).

As schematically shown in Fig. 1, the formation of LAT condensates is initiated by TCR-pMHC binding. A series of kinetic proofreading steps (see below) can lead to the phosphorylation of tyrosine residues on the cytoplasmic tail of LAT and, subsequently, to condensation through a crosslinking reaction with cytosolic crosslinkers. We now elaborate on this process in more detail. When the TCR-pMHC bond life time is sufficiently long (***Stone et al., 2009***; ***Chakraborty and Weiss, 2014***), the binding step is followed by phosphorylation of the ITAMs on the cytoplasmatic tails of the CD3 co-receptors by Lck (***Straus and Weiss, 1992***; ***Van Oers et al., 1996***; ***Lo et al., 2018***). Subsequently, ZAP-70 binds to the ITAM sites and is phosphorylated by Lck as well (***Van Oers et al., 1996***; ***Yan et al., 2013***). In turn, ZAP-70 can phosphorylate the four distal tyrosine residues of LAT (Y132, Y171, Y191, Y226) (***Zhang et al., 1998***; ***Paz et al., 2001***). Phosphorylated tyrosine residues on different LAT molecules can become crosslinked through multivalent interactions with various adapter and signaling proteins including Grb2, SOS1, Gads, SLP76, and PLC*γ*1 (***Zhang et al., 2000***; ***Houtman et al., 2004, Houtman et al., 2006***; ***Zeng et al., 2021***), leading to the formation of LAT clusters and, eventually, LAT condensation. These condensates play a gating role for the propagation of signals downstream of the TCR to Ras/MAPK and calcium pathways (***Samelson, 2002***; ***Huang et al., 2019***; ***McAffee et al., 2022***; ***Morita et al., 2024***; ***Lee et al., 2024***).

**Figure 1.**
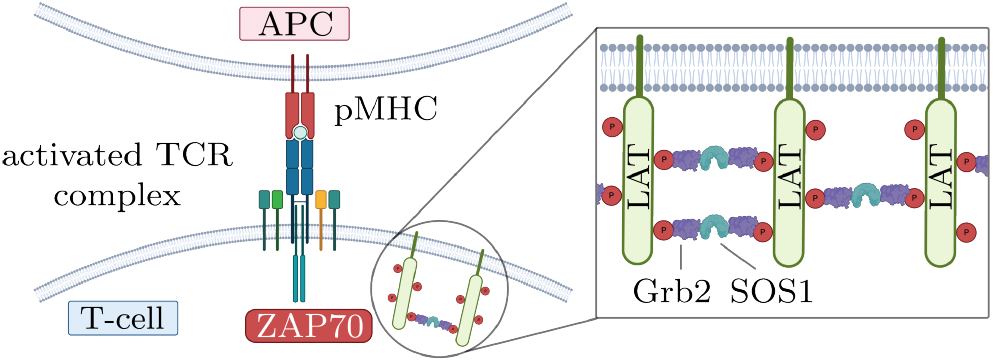
Schematic representation of a T cell engaging with an antigen-presenting cell (APC) through the T-cell receptor (TCR) binding to peptide–MHC (pMHC). Successful recognition of a foreign peptide triggers a signaling cascade that includes the formation of LAT condensates with the help of Grb2 and SOS1 as crosslinkers (inset). LAT condensates serve as hubs that incorporate various other signaling molecules.

Studying the formation of LAT condensates in living cells is challenging, motivating reconstitution experiments on supported lipid bilayers (SLBs) as minimal model systems (***Huang et al., 2016***; ***Su et al., 2016***; ***Huang et al., 2017b***, Huang et al., 2017a; ***Ditlev et al., 2019***; ***Zeng et al., 2021***; ***Sun et al., 2022***). In these reconstitution experiments, the cytoplasmic tail of LAT is attached to a SLB with an N-terminal Histag, as shown in Fig. 2a). In contrast to the physiological system, reconstitution experiments are commonly conducted in the absence of phosphatases such that LAT condensation can be studied at constant phosphorylation levels. Condensation is then initiated by adding the crosslinkers Grb2 and SOS1 to the bulk fluid above the SLB. Thus, reconstitution experiments allow investigating LAT condensates in isolation without spurious effects arising from the complex cellular environment, and have proven highly useful to understand many aspects of the physiological system.

**Figure 2.**
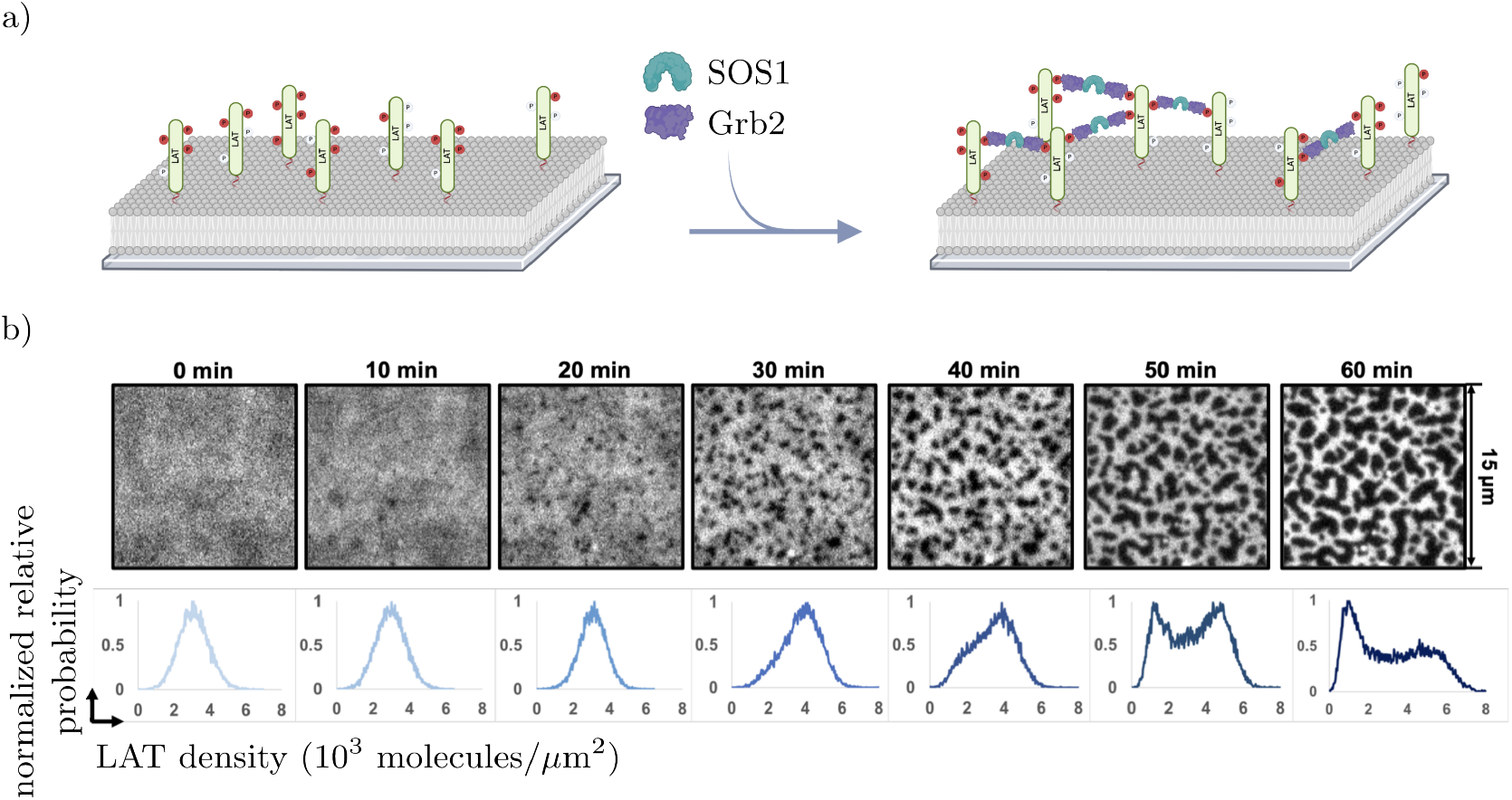
Reconstitution experiments on supported lipid bilayers (SLB). Subfigure a) shows the reconstitution experiment setup. LAT is attached to the SLB via a His-tag and pre-phosphorylated. Grb2 and SOS1 are added to the fluid domain above the SLB, triggering crosslinking and phase separation (Created with Biorender.com). Subfigure b) shows an experimental trajectory where LAT is fluorescently labeled. After an initial lag time without macroscopic density changes, LAT quickly phase separates, leading to a LAT-rich phase with LAT-depleted, non-circular inclusions. This behavior is confirmed by the relative probability of the LAT density, which changes from a distribution with a single peak in the homogeneous phase to a bimodal distribution upon phase separation. The figure was created from data described by ***Sun et al. (2022***).

In these reconstitution experiments, LAT undergoes phase separation into LAT-rich and LAT-depleted micrometer-scale regions that can be imaged by total internal reflection fluorescence microscopy. One realization of a reconstitution experiment is presented in Fig. 2b), with LAT fluorescently labeled, showing an initial time when LAT appears to remain uniformly distributed, followed by rapid phase separation into LAT-depleted domains in a LAT-rich phase. This observation is also reflected in the emergence of a bimodal probability distribution of the LAT density. While previous modeling work provided valuable insights into these experiments (***Nag et al., 2009, Nag et al., 2012***; ***Sun et al., 2022***), we aim to deepen our understanding of the formation of LAT condensates reconstitution experiments by addressing open questions discussed in the following paragraphs. Before proceeding, we note that the following features are also similarly observed in analogous reconstitution experiments with Epidermal Growth Factor Receptor (EGFR) (***Lin et al., 2022b***; ***Phan et al., 2024***).

Reconstitution experiments of LAT condensate formation exhibit a number of intriguing features: Upon the addition of Grb2 and SOS1, we observe an initial lag time of up to 30 min where the LAT density remains homogeneous on length scales greater than about 1 *μ*m, limited by experimental resolution^1^ (see Fig. 2b)) (***Huang et al., 2016***; ***Sun et al., 2022***). However, once phase separation is detected, it proceeds rapidly until phase separation arrests for the remainder of the experiment, resulting in near-stationary patterns undergoing only minor fluctuations (see Fig. 2 and Movie S1).

The observed patterns originating from LAT condensation can be broadly categorized into three distinct morphology groups. The first group consists of a bicontinuous phase where both the LAT-rich and the LAT-depleted phases are interlaced (Fig. 3, left panel). In this case, either the LAT-rich or the depleted phases could make up the majority phase by area fraction. The second and third groups are composed of the minority phase embedded in a percolated majority phase. However, they are distinguished by the geometry of the minority phase, which can be either droplet-like and circular (Fig. 3, middle panel) or elongated and non-circular (Fig. 3, right panel).

**Figure 3.**
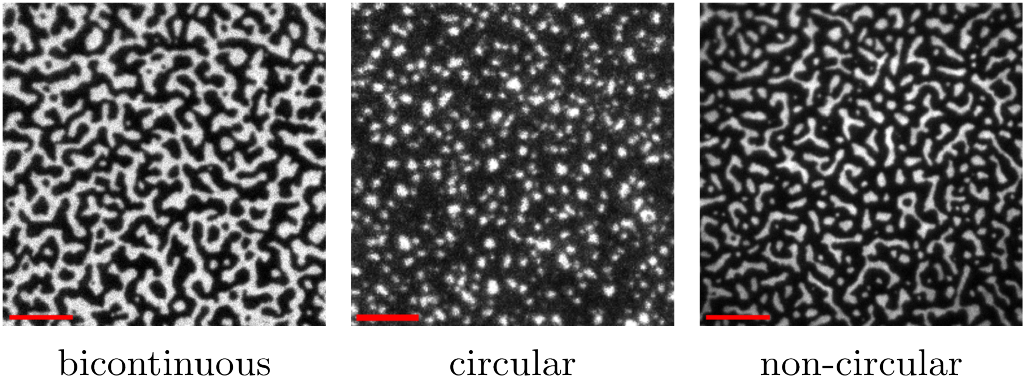
Different morphologies observed in reconstitution experiments, where LAT is fluorescently labeled. The LAT-rich minority phase can form circular or non-circular patterns and similar morphologies can be observed for a LAT-depleted minority phase (see Fig. 11). In addition, the experiments can result in a bicontinuous phase. The figure was created from data described by ***Sun et al. (2022***). (scale bars: 5 *μ*m)

The above observations lead to four major questions we aim to address in this article to gain a better understanding of LAT condensate formation in reconstitution experiments. (1) The characteristic time scale of diffusion of monomeric LAT is significantly shorter than the observed lag time (***Huang et al., 2017a***). Thus, we seek to understand the emergence of this longer lag timescale. (2) What is the origin of the different LAT condensate morphologies shown in Fig. 3. (3) Once condensate formation is initiated, the observed morphologies quickly reach an apparent steady-state. We will try to elucidate this rapid formation of stationary patterns. (4) Albeit not commonly reported, LAT condensate formation can also fail altogether. However, it remains unknown which experimental conditions could prohibit the onset of phase separation.

To address the aforementioned questions, the remainder of this article proceeds as follows. We begin by introducing the Smoluchowski aggregation model together with the aggregation kernels derived by ***Moncho-Jordá et al. (2001***). Using this model, we then identify the mechanisms that give rise to the time delay observed in experimental studies. The aggregation model further motivates the construction of a field-theoretic model that resolves the spatiotemporal dynamics of LAT condensation and thus provides insight into the questions noted above regarding the LAT condensate morphologies observed in experiments.

### Smoluchowski Aggregation Model

To answer the first of the four questions posed in the introduction regarding the origin of the lag time, we employ the Smoluchowski aggregation model (***Smoluchowski, 1916***). The Smoluchowski aggregation model was originally devised for the study of coagulation of colloidal particles but has since been applied to model various other processes such as flocculation of biological cells (***Han et al., 2003***), reversible polymerization (***Van Dongen and Ernst, 1984***), and rain shower onset (***Pruppacher et al., 1998***). In this article, we use the Smoluchowski aggregation model to probe the change of the average cluster size of LAT in time.

#### Model

The Smoluchowski aggregation model describes the time evolution of the concentration of particle aggregates under well-mixed conditions. For LAT condensates in reconstitution experiments, aggregation is effected by the crosslinking of LAT molecules via Grb2 and SOS1. To formulate the model, we consider clusters containing *i* LAT molecules with number density *c*_*i*_(*t*) on the SLB. Two clusters of sizes *i* and *j* can bind with the rate *k*_*ij*_ to form a new cluster of size *i* + *j*. By only accounting for two-body binding events, the Smoluchowski aggregation model is written as

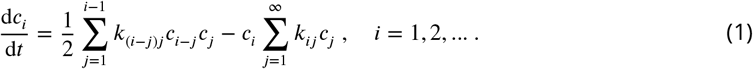

The first term on the right-hand side of Eq. 1 describes the formation of a cluster of size *i* through binding of clusters of sizes *i* − *j* and *j* while the second term captures elimination of clusters of size *i* through the binding to a cluster of size *j*. To solve Eq. 1, we impose the initial condition that only monomers are present at *t* = 0,

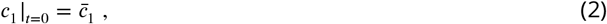

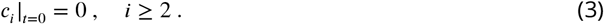

Note that Eq. (1) describes irreversible aggregation, i.e. the fragmentation of clusters is not accounted for. We will discuss the role of fragmentation for the formation of LAT condensates at the end of this section.

The coupled system of differential equations in Eq. (1) needs to be closed by prescribing the form of the rate constants *k*_*ij*_. To obtain expressions for these rate constants, the limits of diffusion-limited cluster aggregation (DLCA) and reaction-limited cluster aggregation (RLCA) are commonly considered (***Müller, 1926***; ***Jullien, 1992***). DLCA is marked by high binding affinities and slow diffusion while RLCA results from low binding affinities and fast diffusion. From the diffusion coefficient of monomeric LAT, we find that diffusion is initially fast compared to the experimental time scale (***Huang et al., 2017a***), suggesting RLCA is the dominant aggregation mechanism. However, as clusters grow, diffusion slows down and we expect a transition from RLCA to DLCA. To capture this transition, the rate constants *k*_*ij*_ are split into Brownian and reactive contributions (***Moncho-Jordá et al., 2001***),

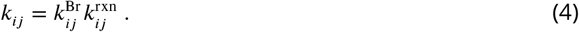

The role of 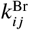 and 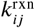 is depicted in Fig. 4. The Brownian contribution 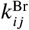 describes the rate at which clusters of size *i* and *j* come within a distance where they could bind, referred to as an encounter (***Moncho-Jordá et al., 2001***), and 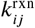 describes the probability that such an encounter leads to binding of the two clusters.

**Figure 4.**
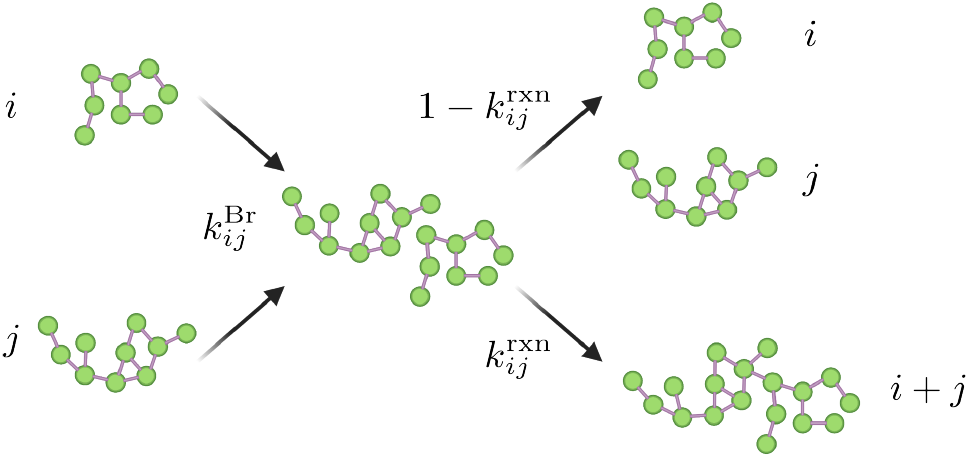
Illustration of the multiplicative split of the reaction kernels proposed by ***Moncho-Jordá et al. (2001***). The Brownian contribution 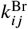 provides the rate at which clusters of size *i* and *j* get within binding distance and 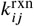 describes the probably of the clusters to subsequently bind during this encounter (Created with Biorender.com)

In choosing the form of the Brownian and reactive contributions, we follow the rate constants proposed by ***Moncho-Jordá et al. (2001***). Specifically, the reactive contribution takes the form

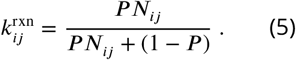

In this expression, *P* is the binding probability of two clusters upon collision and *N*_*ij*_ is the number of collisions per encounter of clusters of size *i* and *j* before they diffuse apart. While the form of *N*_*ij*_ is unknown, it is argued by ***Moncho-Jordá et al. (2001***, Moncho-Jordá et al. 2004) that it must be an increasing function of *i* and *j*. Adopting the ansatz from these references, we use the expression

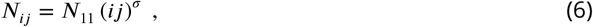

with *N*_11_ denoting the number of collisions per encounter for two monomers and *σ* > 0 being a free parameter.

The Brownian contribution to the binding rate 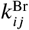 is determined by the diffusion coefficients of the clusters, i.e.

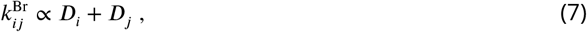

with *D*_*i*_ and *D*_*j*_ denoting the diffusion coefficients of clusters of size *i* and *j*^2^. To proceed, we note that colloidal aggregation through both DLCA and RLCA leads to fractal clusters of particles (***Meakin, 1987b***; ***Robinson and Earnshaw, 1992***). In three-dimensions, the diffusion coefficients of such aggregates can be evaluated using Kirkwood-Riseman theory (***Kirkwood and Riseman, 1948***; ***Meakin et al., 1985***) or the porous sphere model (***Debye and Bueche, 1948***; ***Warren, 1994***). While extensions to the collective diffusion of particles exist for free-standing lipid membranes (***Oppenheimer and Diamant, 2009***; ***Sokolov and Diamant, 2018***; ***Sorkin and Diamant, 2021***), they have not been applied to fractal aggregates or adapted to SLBs. Therefore, we make the ansatz that the diffusion coefficients depend on the the hydrodynamic radius 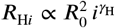, where *R*_0_ is the radius of gyration of a monomer and *γ*_H_ is the mobility-mass scaling exponent^3^(***Sorensen, 2011***). We can then use the Evans-Sackmann theory to relate the hydrodynamic radius to the diffusion coefficient of a cluster containing *i* particles (***Evans and Sackmann, 1988***),

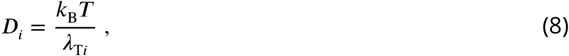

with the translational drag coefficient given by

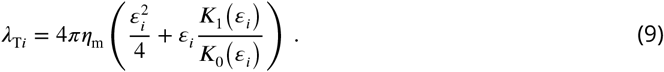

Here, *η*_m_ denotes the membrane viscosity, 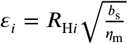 is the effective dimensionless radius with *b*_s_ being the friction coefficient describing the membrane-substrate interactions, and *K*_0_ and *K*_1_ are modified Bessel functions of the second kind. The drag coefficient in Eq. (9) is dominated by the hydrodynamic drag arising from the membrane viscosity at small scales (*D*_*i*_ ∝ − ln *ε*_*i*_) and by the membrane-substrate friction at large scales 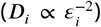. This form of the diffusion coefficient has also been confirmed experimentally (***Kaizuka and Groves, 2004***).

Before proceeding, we recall that the form of the aggregation model in Eq. (1) does not capture the effects of fragmentation of clusters resulting from the unbinding of crosslinkers (***Huang et al., 2016***). Accounting for fragmentation would lead to additional terms in Eq. (1) that are linear in the concentrations *c*_*i*_ (***Krapivsky et al., 2010***). At early times, the concentration of all clusters of size *i* ≥ 2 is small compared to the monomer concentrations, suggesting that the fragmentation of these clusters does not qualitatively change the dynamics even though it could lead to an effective slowing of cluster growth. At later times, we expect that the high valency of LAT quickly results in the bulk energy of a cluster dominating over its interfacial energy, thus stabilizing larger clusters against fragmentation, consistent with classical nucleation theory (***Karthika et al., 2016***). Thus, we do not expect fragmentation of clusters to qualitatively alter the results presented in the next section.

To analyze the Smoluchowski aggregation model, we numerically integrate the system of ODEs given by Eq. (1) with the reaction kernels defined by Eqs. (4)–(9) and subject to the initial conditions in Eqs. (2) and (3). Table 1 shows the parameters used in our results unless stated otherwise. For the friction coefficient *b*_s_ and membrane viscosity *η*_m_, we use typical values found in the literature. However, values for the binding probabilty *P*, collision frequency parameters *N*_11_ and *σ*, and the mobility-mass scaling exponent *γ*_H_ are not known. We focus on varying the parameter *P* in the following and show in the SM (Fig. S1) that the obtained results do not qualitatively depend on the values of the remaining parameters.

**Table 1.** Parameters used to numerically solve the Smoluchowski aggregation model.

|  |  |
| --- | --- |
| $P$ | $10^{-2}$ |
| $N_{11}$ | 1 |
| $\sigma$ | 1 |
| $\eta_m$ | $10^{-1}$ pN $\mu\text{s}/\text{nm}$ (*Honerkamp-Smith et al., 2013*) |
| $b_s$ | $10^{-2}$ pN $\mu\text{s}/(\text{nm})^3$ (*Anthony et al., 2022*) |
| $\gamma_H$ | $2/3$ (*Meakin, 1987a*) |

The numerical integration is implemented using MATLAB’s ode45 solver (***The MathWorks Inc., 2024***). To keep the integration computationally tractable, we solve Eq. (1) for *i* ≤ *N*_max_ = 4000 and terminate our simulations once 5 % of the initial mass has been lost to clusters of size *i* > *N*_max_. In the following, we present our results in terms of the mean cluster size, *s* = Σ_*i*_ *ic*_*i*_/ Σ_*j*_ *c*_*j*_ and the non-dimensional time 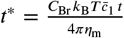, where *C*_Br_ is the proportionality factor in Eq. (7).

## Results and discussion

Figure 5, left panel, shows the mean cluster size for several values of the binding probability *P*. In all cases, we find an initial lag time without any significant cluster growth. This initial lag time increases with decreasing *P*. The lag time is followed by a rapid increase of the average cluster size that appears independent of *P*. After sufficiently long times, the rate at which the average cluster size grows decreases. The time and mean cluster size at which the growth rate decreases also depends on the value of *P*.

**Figure 5.**
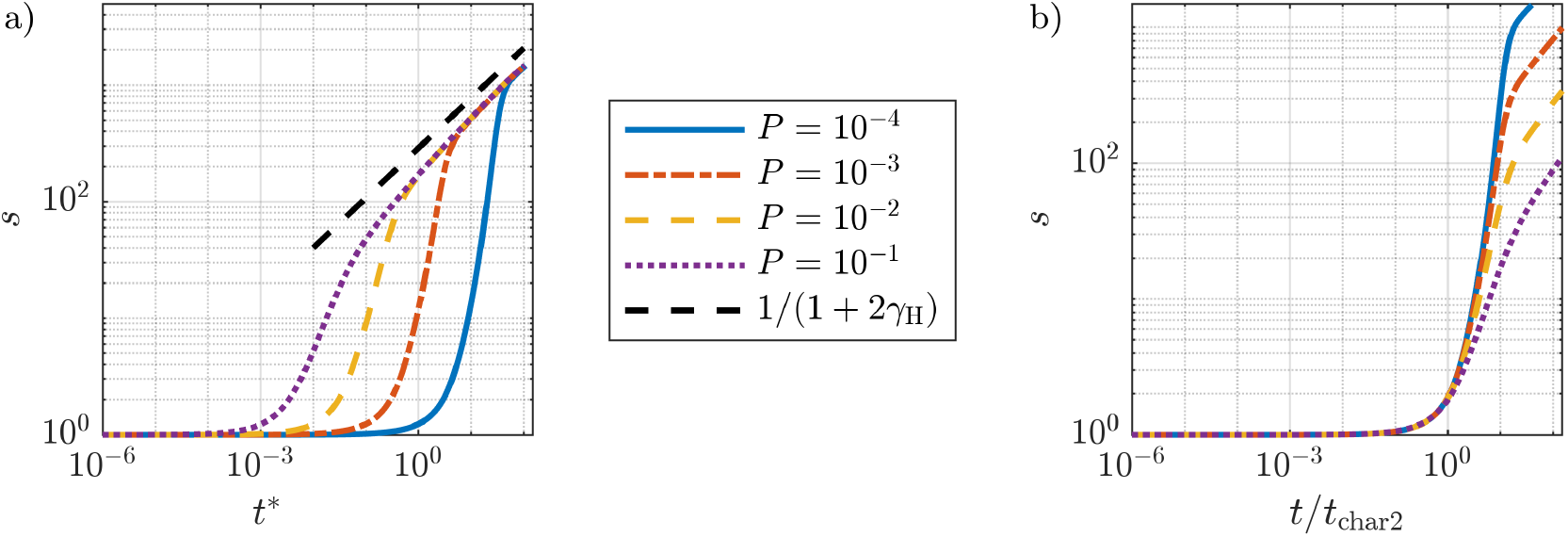
Mean cluster size *s* = Σ_*i*_ *ic*_*i*_/Σ_*j*_ *c*_*j*_ obtained from simulating Eqs. (1)–(9) for different values of the binding probability *P*, plotted against non-dimensional time *t*^*^ (left panel) and time rescaled by the characteristic time of dimer formation, *t*_char2_ (right panel). The onset of a rapid increase in the average cluster size follows an initial lag time that is modulated by *P*. Scaling of time by the characteristic time of dimer formation collapses all curves at early times in the right panel, indicating that the lag time originates from the initial formation of smaller aggregates.

We first seek to understand the origin of the lag time. To this end, consider the concentration of clusters of size *i* > 1. The initial condition in Eq. (3) suggests that the concentrations *c*_*i*_ increase from zero as clusters of size *i* form from smaller clusters. Since we consider irreversible aggregation, clusters of size *i* are continuously converted to larger clusters, suggesting that *c*_*i*_ reaches a maximum and subsequently decreases (see also ***Krapivsky et al. (2010***)). In Sec. 1.1 of the SM, we find an approximate expression for the time at which the dimer concentration is maximum, i.e.

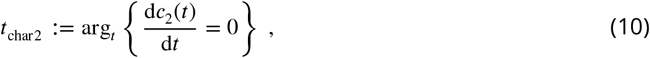

yielding

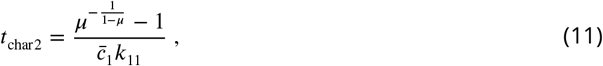

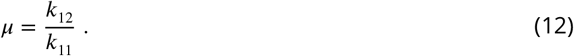

In Fig. 5, right panel, we rescale time by *t*_char2_ and find that the mean cluster size collapses for all values of *P* at early times and that *t*/*t*_char2_ ≈ 1 when the cluster size starts to rapidly increase. This suggests that *t*_char2_ indeed represents the lag time and further indicates that the onset of aggregation of LAT molecules can only proceed after initial formation of small dimers.

To obtain the scaling of the characteristic time with the binding probability *P*, we note that for *P* ≪ 1, *μ* is approximately independent of *P*. From this, we immediately obtain the result

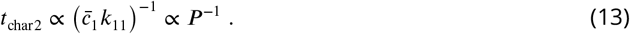

This shows that the scaling of *t*_char2_ is determined by the monomer concentration and the rate at which monomers can be converted to dimers.

For *t* > *t*_char2_, the average cluster size increases rapidly. In this regime, cluster growth is determined by both the reactive and diffusive contributions to *k*_*ij*_ and an expression for the growth rate is currently not known. However, we observe a second crossover to a lower growth rate with the time of this crossover increasing with decreasing *P*. The lower growth rate is associated with the transition to diffusion-limited cluster aggregation. In this regime, the growth rate of *s* follows a power law relation,

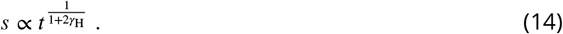

This result can be obtained from the scaling ansatz described by ***Krapivsky et al. (2010***) and the scaling of the diffusion coefficient for large clusters in Eq. (8) as 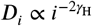. Furthermore, the cluster size at which this transition occurs can be determined by finding *i*_cross_ such that 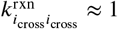, yielding the crossover cluster size for *P* ≪ 1 as (***Moncho-Jordá et al., 2001***)

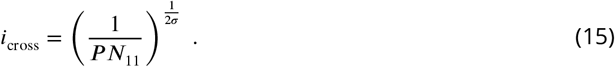

In the SM (Fig. S2), we rescale the mean cluster size by *i*_cross_, showing that the slowing of the growth rate is indeed determined by the transition to DLCA.

In summary, the Smoluchowski aggregation model showed that the lag time can be explained by a low binding probability of LAT monomers. This low binding probability originates from LAT binding with the help of a Grb2 and SOS1 as crosslinkers: Aggregation does not only require two LAT molecules to come within binding distance on the membrane, it also requires the crosslinkers to bind to LAT in the correct stoichiometric ratio, leading to an effectively low binding probability in experiments. Furthermore, the binding probability is affected by the average number of phosphorylated sites, allosteric effects (***Huang et al., 2016***) and the binding of monomeric Grb2 or Grb2:SOS1 dimers to LAT instead of the required Grb2:SOS1:Grb2 trimer (***Huang et al., 2016***), effectively blocking tyrosine residues. Lastly, we discussed above that the model in Eq. (1) assumes binding to be irreversible. If reversibility is considered, however, the onset time would likely not be governed by the formation of dimers but larger multimers that are sufficiently stable against fragmentation. Thus, we expect our results to remain qualitatively valid if fragmentation of dimers and small clusters is taken into account.

### Field-Theoretic Model

The Smoluchowski aggregation model discussed in the previous section revealed that the onset of rapid cluster growth can be explained by a low effective binding probability. This result clarifies the question posed the Introduction section regarding the origin of the lag time observed in experiments. To address questions (2)–(4) noted in the Introduction, we propose a field-theoretic model that spatially resolves the dynamics of LAT and thus captures the experimentally observed morphologies after phase separation.

#### Model

To construct the field-theoretic model, we employ the insights obtained from the Smoluchowski aggregation model. Specifically, we found that we require sufficiently many dimers to form before nucleating the growth of large clusters. As discussed in the Smoluchowski Aggregation Model section, a threshold number of small clusters instead of dimers is needed in the case of reversible binding. This is equivalent to requiring a sufficiently large average number of bonds per LAT molecule. Once sufficiently many bonds have formed, cluster growth proceeds rapidly and macroscopic phase separation is observed. To incorporate the above idea into a field-theoretic model, we introduce the LAT density, *L*(***x***, *t*), and the bond density per LAT molecule, *ϕ*(***x***, *t*) *∈* [0, *ν*/2], where *ν* is the average valency per LAT molecule. To model the evolution of the two fields, we propose two coupled dynamical equations.

Since LAT is conserved, the evolution of *L*(***x***, *t*) can be related to a flux ***j***_*L*_[*ϕ*(***x***, *t*), *L*(***x***, *t*)], as

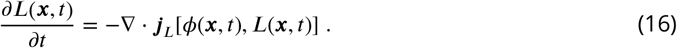

We assume that the flux ***j***_*L*_[*ϕ*(***x***, *t*), *L*(***x***, *t*)] can be obtained as the gradient of a free energy functional ℱ[*ϕ*(***x***, *t*), *L*(***x***, *t*)] (***Hohenberg and Halperin, 1977***), but with a mobility that also depends on both densities, i.e.

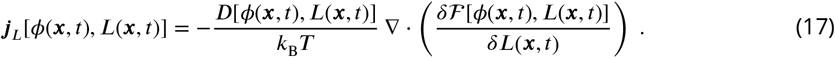

To construct the free energy functional, we employ an analytic expansion that is linear in *ϕ* and quartic in *L*. This is the lowest-order model exhibiting phase separation regulated by the bond density *ϕ*. With this choice, ℱ[*ϕ*(***x***, *t*), *L*(***x***, *t*)] takes the generic form

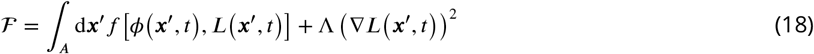

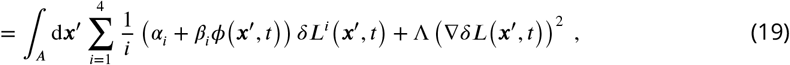

where *f* [*ϕ*(***x***, *t*), *L*(***x***, *t*)] is the *bulk area* free energy density, Λ denotes the line tension, 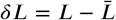 is the LAT density with respect to some constant reference value 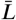 and *α*_*i*_ and *β*_*i*_ are parameters of the free energy density. To reduce the number of parameters, we postulate that the free energy should take the characteristic form shown in Fig. 6a), where the red dots indicate the minima of the free energy density *f* at fixed *ϕ*. Specifically, we require that there is only one minimum for *ϕ* ≤ *ϕ*_th_, for some threshold value of the bond density, *ϕ*_th_, and two minima for *ϕ* > *ϕ*_th_. As shown in Sec. 2.1 of the SM, enforcing these constraints reduces the permissible free energy density to

**Figure 6.**
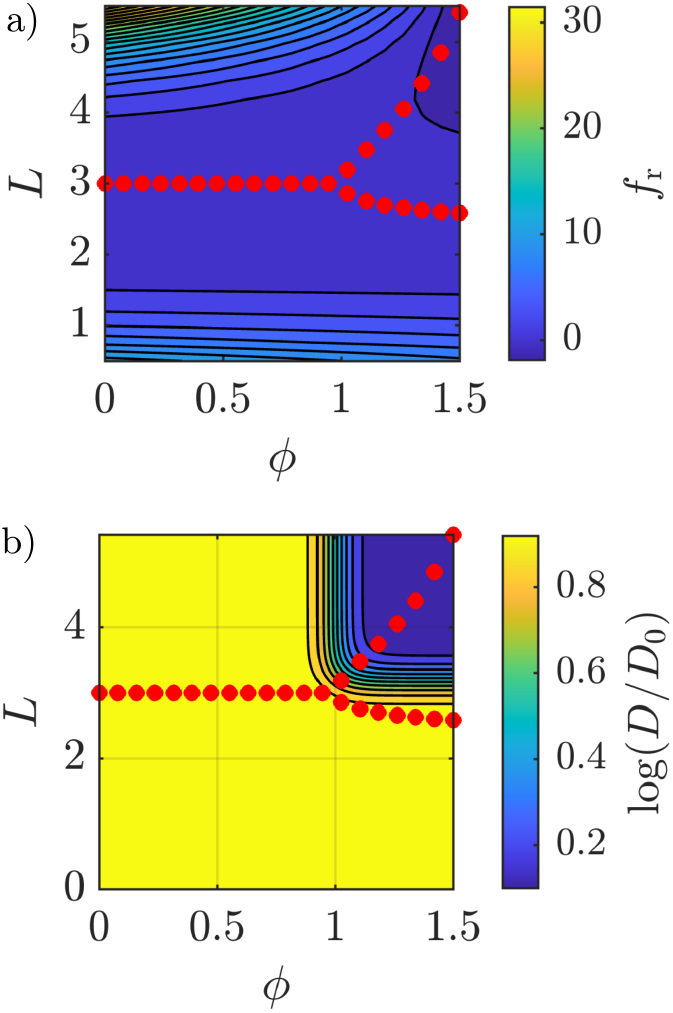
Free energy density a) and diffusion coefficient b) for *ϕ*_th_ = 1 and 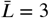. The red dots indicate the minima of the free energy density *f*_r_. Subfigure a) shows that only a single minimum in the energy density exists for *ϕ* < *ϕ*_th_ and two distinct minima for *ϕ* > *ϕ*_th_. If the LAT density is close to the minimum with the higher LAT density, the diffusivity is significantly reduced, as can be seen in Subfig. b). Otherwise, the diffusivity remains unaffected.

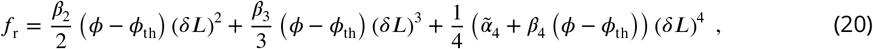

with the new parameter 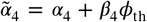. In addition, the parameters in Eq. (20) are subject to the constraints

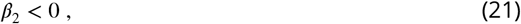

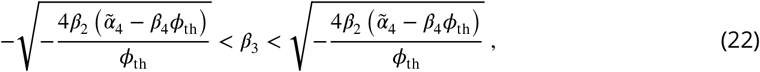

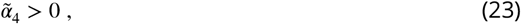

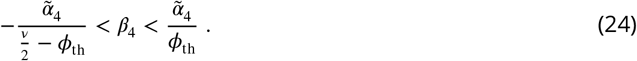

Substituting Eqs. (18) and (20) into Eq. (17) yields the flux of LAT as

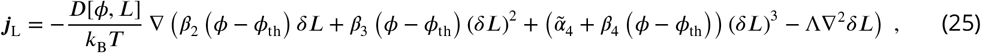

and using this result in Eq. (16) leads to the evolution equation for *L*(***x***, *t*),

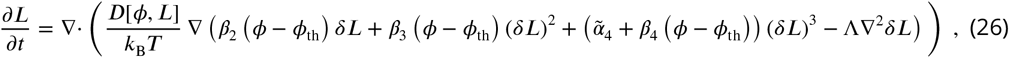

where we have now omitted the explicit dependencies on space and time for clarity.

In the context of the Smolchowski aggregation model, we discussed that the diffusion coefficient decreases with the cluster size and used the Evans-Sackmann theory to quantify this. This effect has also been experimentally measured in the LAT-rich phase (***Huang et al., 2017a***), motivating the dependence of the diffusion coefficient on the bond and LAT densities in Eq. (17). However, in the field-theoretic model, there does not exist a measure of the size of an aggregate such that the Evans-Sackmann theory cannot be applied. Therefore, we propose a phenomenological model of the diffusion coefficient,

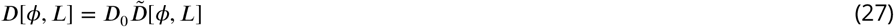

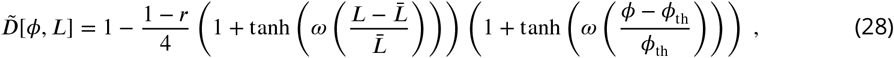

with *α* describing the change in diffusivity, and *ω* being the width of the transition region where the diffusivity changes from *D*_0_ to *D*_0_*α*. An example of Eq. (28) is shown in Fig. 6b). We note that the dependence of the expression in Eq. (28) on 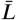 and *ϕ*_th_ ensures that the diffusion coefficient is not decreased before phase separation occurs, i.e. 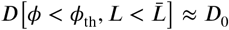.

Equations (16)–(28) establish the governing equation for the LAT density given the bond density per LAT *ϕ*. We propose that the dynamics of *ϕ* follow a reaction-diffusion equation,

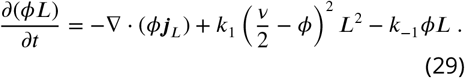

This equation is expressed in terms of the bond density *ϕL* instead of the bond density per LAT to allow for an easier interpretation. The first term on the right-hand side of Eq. (29) describes that a diffusing LAT molecule carries *ϕ* bonds along with it, leading to the flux of bonds *ϕ****j***_*L*_. The remaining two terms describe the binding of two LAT molecules via a crosslinker from the bulk fluid and the corresponding unbinding reaction. We note that the rate constants *k*_1_ and *k*_−1_ also describe the effects of the bulk concentrations of Grb2 and SOS1 on the binding propensity of LAT. In here, we assume that the bulk fluids act as reservoirs for Grb2 and SOS1 such that we can assume *k*_1_ and *k*_−1_ to be constant in time.

To solve the field-theoretic model given by Eqs. (26) and (29), we derive their non-dimensional forms in Sec. 2.2 of the SM. Using *asterisks* and *hats* to describe non-dimensional fields and parameters, respectively, this yields the non-dimensional flux

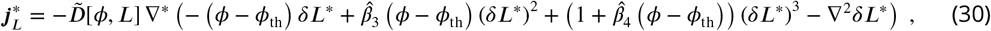

and the dynamical equations

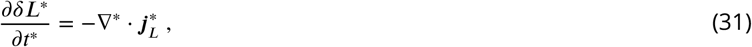

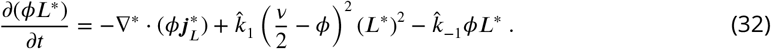

The non-dimensional parameters in Eqs. (30)–(32) are given by

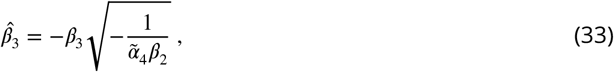

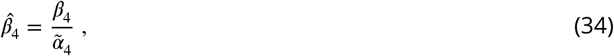

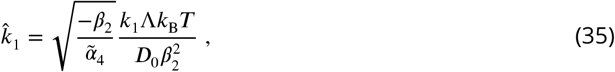

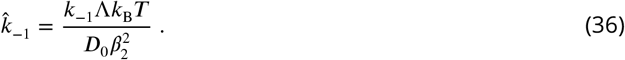

For ease of notation, we will omit the *asterisk* symbol in the following.

We solve Eqs. (30)–(32) on a square domain subject to periodic boundary conditions using the Dedalus Project (***Burns et al., 2020***), a python package implementing pseudospectral methods for partial differential equations. Using a Fourier basis and Dedalus’ 2nd-order Crank-Nicolson Adams-Bashforth (*CNAB2*) time stepper, we evolve the dynamical equations subject to the initial conditions,

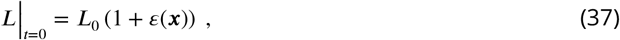

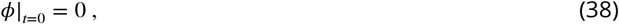

where *L*_0_ is the average initial LAT density and *ε*(***x***) ≪ 1 is a spatially varying, uniformly distributed noise term with zero mean. We run the spatially-resolved simulations for a fixed time *T*_sim_ (see Tab. 2) and record the observed patterns at regular intervals. For the phase diagrams in Figs. 10 and 12, we only consider the final recorded snapshots of the simulations.

**Table 2.** Parameter values used in Eqs. (30)–(32) for the spatially-resolved simulation results.

| $\bar{L}$ | $\omega$ | $r$ | $\nu$ | $\phi_{\text{th}}$ | $\hat{k}_1$ | $\hat{k}_{-1}$ | $T_{\text{sim}}$ |
| --- | --- | --- | --- | --- | --- | --- | --- |
| 3 | 20 | 0.05 | 3 | 1 | $\frac{1}{3} \times 10^{-4}$ | $\frac{1}{3} \times 10^{-5}$ | $6 \times 10^4$ |

## Results and discussion

In this section, we present the results obtained by solving the model proposed given by Eqs. (30)–(32). We begin by considering the time before phase separation can occur, i.e. when *ϕ* < *ϕ*_th_. During this time, the diffusive flux ***j***_*L*_ is small and the LAT density remains approximately homogeneous. We can then solve Eq. (32) to determine the time when *ϕ* reaches *ϕ*_th_. In the flux-free case, Eq. (32) is the Riccati equation, which can be cast as a linear second-order differential equation. By solving this differential equation subject to Eq. (38), we obtain

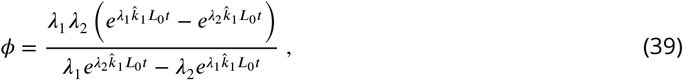

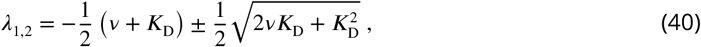

where we introduced the dissociation constant

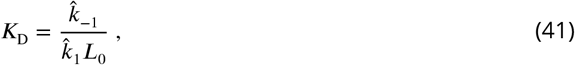

and assumed *K*_D_ > 0. Figure 7a plots the bond density *ϕ* against the non-dimensional time 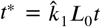 for different values of the dissociation constant *K*_D_ according to Eq. (39). At early times, the majority of binding sites are vacant, leading to a rapid increase in bond density. As the bond density increases, the number of available sites decreases, while unbinding additionally counters binding. This effects a slower rate of bond formation until binding and unbinding are balanced and a steady-state is reached. The steady-state bond density decreases with increasing dissociation constants.

**Figure 7.**
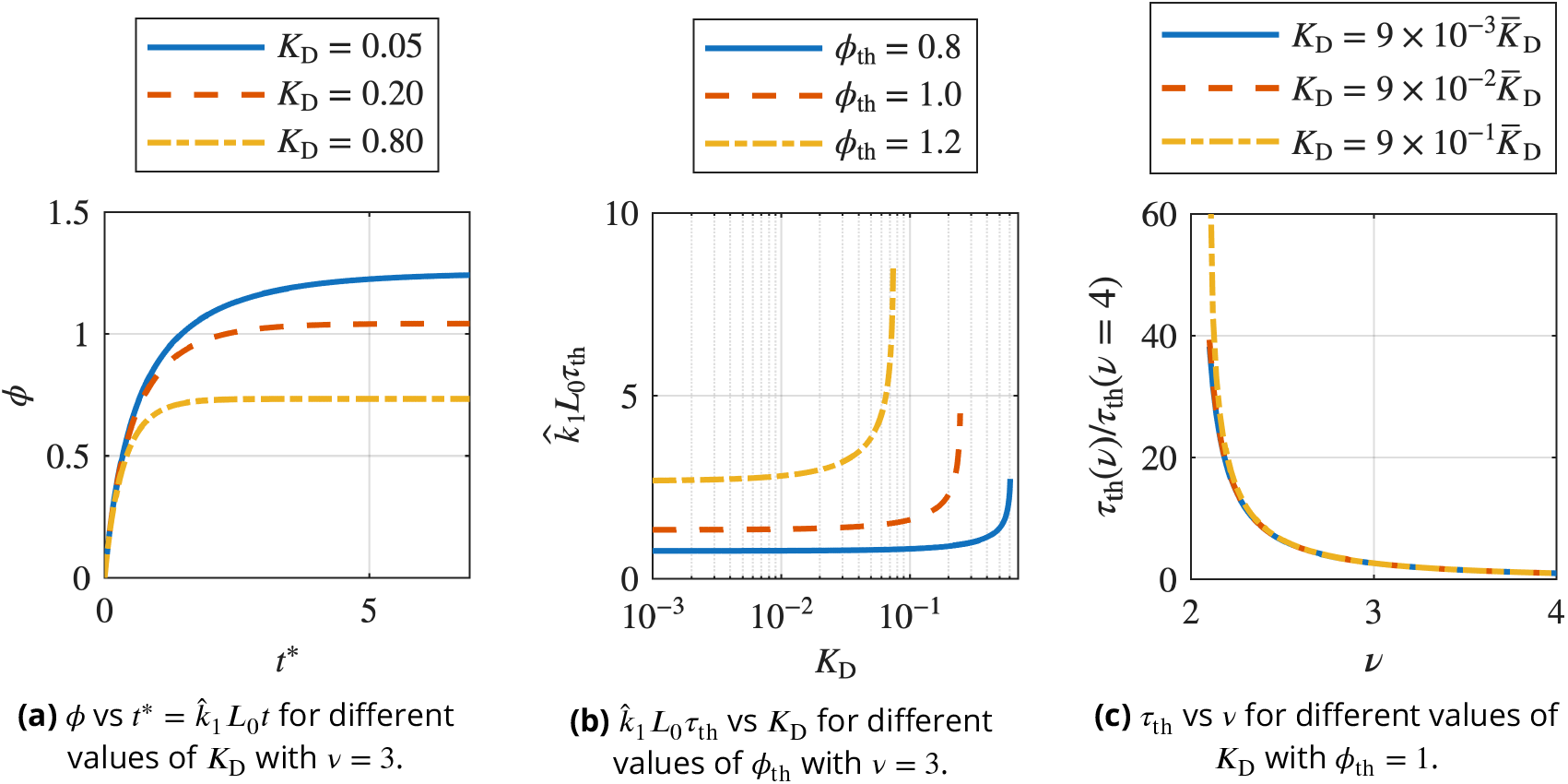
Bond formation in the homogeneous state before phase separation. Subfigure (a) shows the bond density plotted against non-dimensional time for different values of the dissociation constant *K*_D_. The bond density increases rapidly initially and subsequently plateaus at a steady-state value that decreases with increasing *K*_D_. Subfigure (b) shows the time *τ*_th_ to reach *ϕ* = *ϕ*_th_, i.e. the threshold bond density required for phase separation. When *K*_D_ approaches 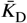, defined in Eq. (42), *τ*_th_ increases rapidly as the effects of bond dissociation become relevant. Lastly, Subfig. (c) shows *τ*_th_ for *ϕ*_th_ = 1 depending on the average number of binding sites per LAT molecule normalized by the case *ν* = 4 for each value of *K*_D_. We observe a several-fold increase of *τ*_th_ as the number of available binding sites decreases.

The dependence of the steady-state bond density on the dissociation constant suggests there exists a maximum value of *K*_D_, denoted by 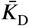, such that the threshold bond density *ϕ*_th_ cannot be reached for 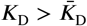. Indeed, we find that the dissociation constant *K*_D_ must satisfy

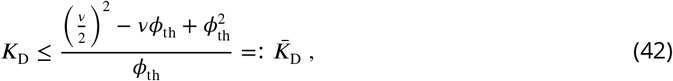

for *ϕ* to exceed *ϕ*_th_ as *t* → ∞. Otherwise, *ϕ* equilibrates at a value *ϕ* < *ϕ*_th_ and phase separation can not occur. If Eq. (42) is satisfied, we obtain the time required to reach *ϕ*_th_ from Eq. (39) as

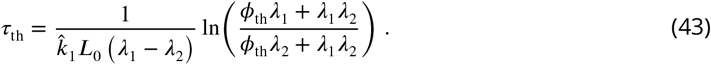

In Fig. 7b, the non-dimensional time 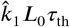 is plotted against the dissociation constant for different values of the threshold bond density *ϕ*_th_, commonly assumed to satisfy *ϕ*_th_ ≈ 1 (***Barisas, 2003***; ***Nag et al., 2009, Nag et al., 2012***). For decreasing values of *K*_D_, the non-dimensional time 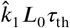 approaches a constant value. Thus, 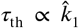, implying that *τ*_th_ is limited by the forward rate of the reaction for small *K*_D_. As *K*_D_ approaches 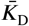 from below, *τ*_th_ increases rapidly due to the increasing rate of unbinding. This shows that the onset of phase separation is modulated by the effective binding constants 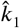 and 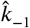 as well as the initial LAT density (cf. Eq. (41)), which is consistent with the Smoluchowski aggregation model (cf. Eq. (13)) while providing additional insight into the lag times observed in experiments.

Lastly, we plot the change of *τ*_th_ as a function of the average number of binding sites per LAT molecule for *ϕ*_th_ = 1 in Fig. 7c. For every value of *K*_D_, *τ*_th_ is normalized by *τ*_th_ when the number of binding sites is maximum, i.e. *ν* = 4. Here, we find that reducing the number of available binding sites can increase *τ*_th_ several-fold. Therefore, an additional time delay can occur when LAT is not efficiently phosphorylated, effectively reducing the number of binding sites. This could, for example, result from phosphatase pressure leading to dephosphorylation of tyrosine residues.

Thus far, we examined the behavior of LAT prior to the onset of phase separation. We now turn to analyzing the dynamics that govern phase separation. However, while our aim is to construct a minimal model, it does involve a number of (non-dimensional) parameters. Namely, the initial LAT concentration *L*_0_, the parameters describing the diffusivity *ω* and *α*, the average number of binding sites per molecule *ν*, the threshold bond density *ϕ*_th_, the free energy parameters 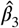 and 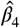 as well as the rate constants 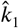 and 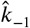. This relatively large number of parameters makes it infeasible to fully explore the parameter space in this article. Therefore, we focus on investigating the effects of the free energy parameters 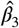 and 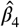 and the initial LAT density *L*_0_. The values of the remaining parameters are listed in Tab. 2. We note here that the value of *α* is larger than measured in experiments (***Huang et al., 2017a***) but of the same order of magnitude. However, smaller values of *α* lead to numerical instabilities in our model, motivating the choice in Tab. 2.

Figure 8 shows an example trajectory obtained from our simulations with snapshots taken at evenly distributed times. In the top row, the colorcoding indicates the LAT density *L* while in the bottom row, the colorcoding indicates the bond density per LAT *ϕ*. At early times, both the LAT and bond density remain homogeneous while the bond density increases due to binding of Grb2 and SOS1 from the bulk fluid. Once the the bond density reaches *ϕ*_th_ between the second and third frame, we observe the onset of phase separation and LAT-rich islands form in a LAT-depleted majority phase. As a result of the reduced diffusion coefficient in the LAT-rich phase, the observed morphology remains near-stationary, as also observed in experiments. However, the density of LAT increases between the final two frames in the LAT-rich phase. Furthermore, we observe that the bond density per LAT molecule is larger in the LAT-rich domains. This is a result of the quadratic dependence on the LAT density in the binding term of Eq. (29).

**Figure 8.**
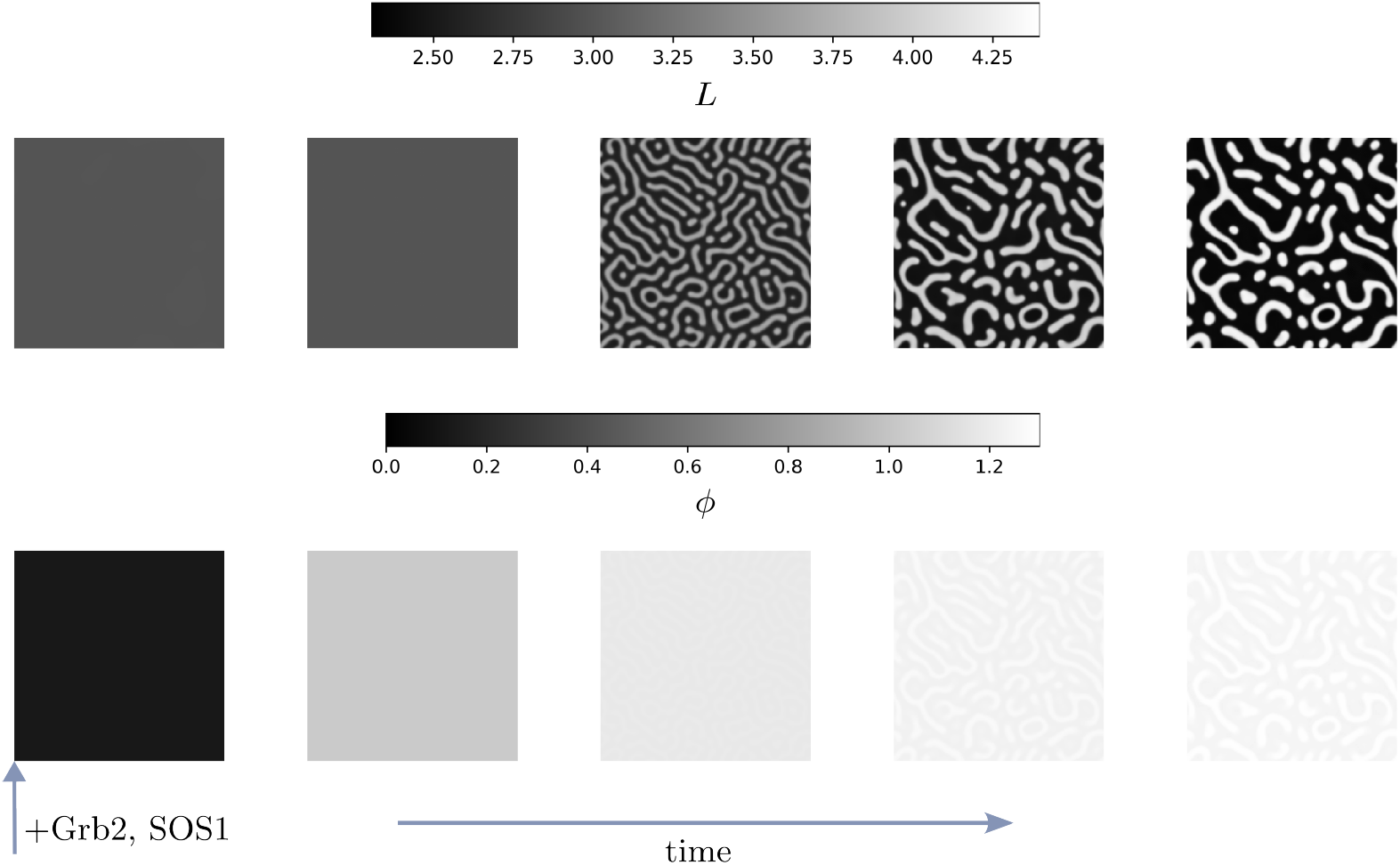
Simulation trajectory obtained for the parameters *L*_0_ = 3.0, 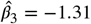 and 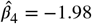. The top row shows the LAT density *L* while the bottom row shows the bond density *ϕ*. At early times, the bond density is small and no phase separation is observed, Between the second and third frame, the threshold bond density is reached and phase separation is observed. Subsequently, elongated, LAT-rich inclusions form in the LAT-depleted majority phase. Due to the small diffusion coefficient in the LAT-rich phase, these inclusions become close to stationary. Furthermore, the non-linear dependence on the binding contribution in Eq. (29) leads to preferred binding in the LAT-rich phase, resulting in an increased bond density in that phase.

While Fig. 8 shows an example trajectory for a choice of parameters that leads to LAT-rich, non-circular domains in a LAT-depleted majority phase, our proposed model can yield all other experimentally observed morphologies. Figure 9 shows the remaining morphologies, namely near-circular LAT-rich and LAT-depleted droplets, non-circular LAT-depleted inclusions as well as a bicontinuous phase.

**Figure 9.**
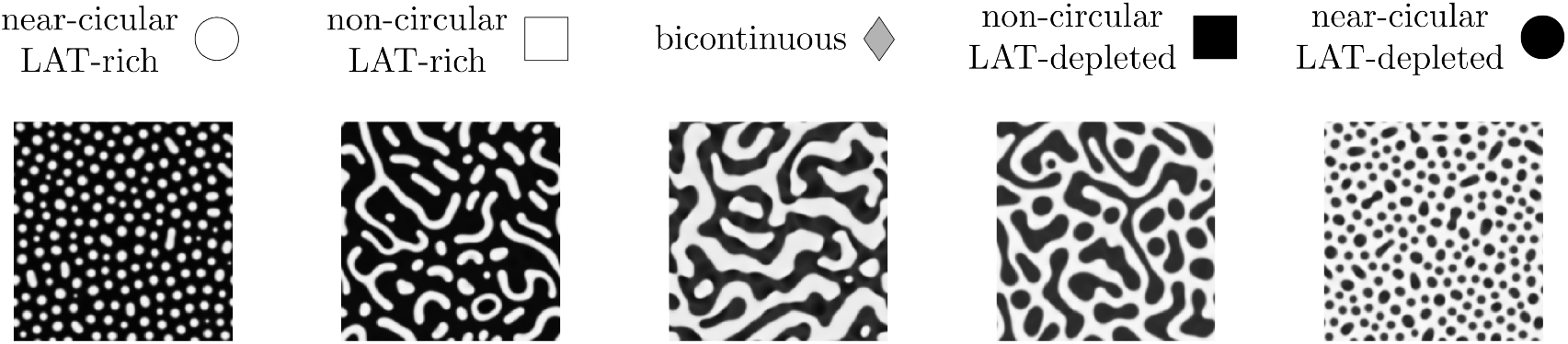
Examples of the different possible morphologies obtained from our simulations, closely resembling the morphologies obtained in experiments.

**Figure 10.**
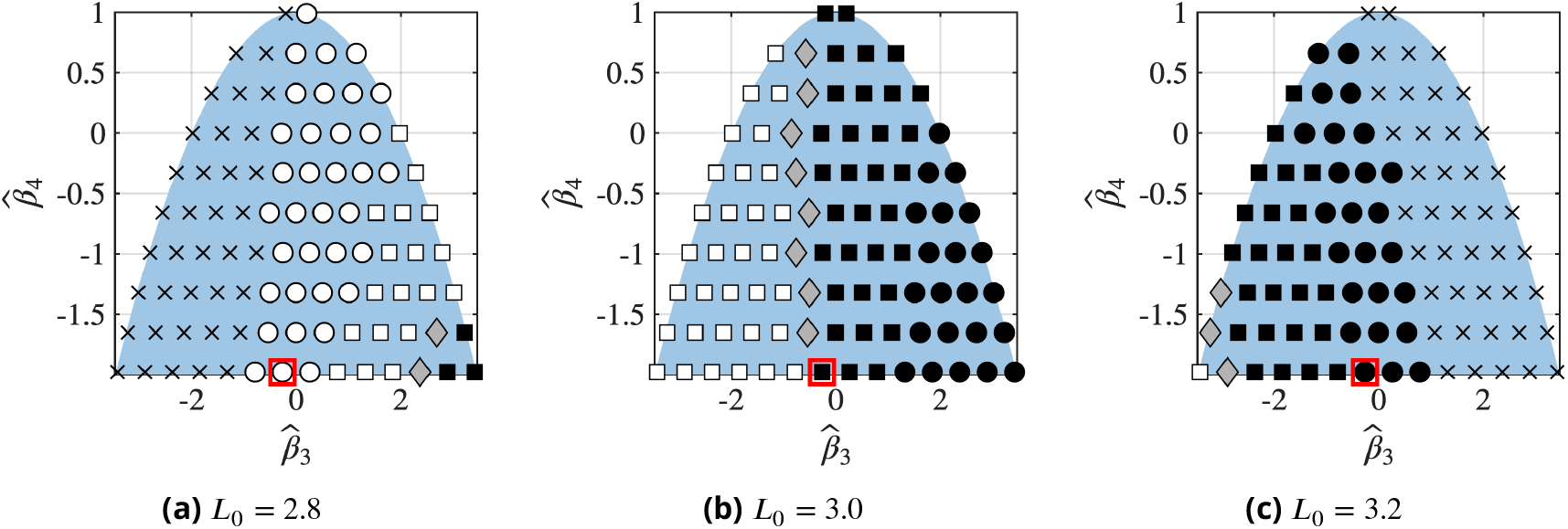
Phase diagrams of the observed morphologies at the end of each simulation for varying free energy parameters 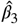 and 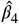, defined in Eqs. (33) and (34), respectively, at different initial densities *L*_0_. The blue background indicates the domain of permissible parameter combinations based on Eqs. (22) and (24). Each marker represents a single simulation and the meaning of each marker is indicated in Fig. 9.

To understand how different parameters affect the resulting morphologies, we systematically varied the non-dimensional free-energy parameters 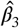 and 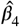, the parameters associated with the cubic and quartic terms in the free energy in Eq. 20, respectively. The results are shown in Fig. 10 for different initial LAT densities *L*_0_. Here, the blue background indicates the permissible parameter range based on the constraints posed by Eqs. (22) and (24). We characterize each simulation result based on the final time step and indicate the morphology using the symbols in Fig. 9. In addition, we use a cross (×) symbol to indicate that no phase separation is observed by the end of the simulation.

We begin by considering Fig. 10b where the initial LAT density is *L*_0_ = 3.0. In this case, phase separation is observed for all admissible parameter combinations. Furthermore, for all values of 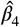, small values of 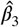 lead to LAT-rich inclusions whereas large values of 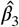 give rise to LAT-depleted inclusions. This is a result of the change of location of the smallest free energy minimum at some *δL* < 0 for 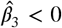 to some *δL* < 0 for 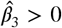. In between these two regimes, we observe a narrow band with a bicontinuous phase. With increasing values of 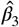, we also observe the transition from non-circular LAT-depleted inclusions to circular LAT-depleted inclusions. This is a result of the free energy minimum associated with the LAT-depleted phase shifting towards lower densities, consequently forming smaller inclusions due to mass conservation. For these smaller inclusions, line tension becomes dominant, driving them towards circular shapes.

An analogous transition to circular LAT-rich inclusions is not observed for 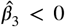, despite the form of the free energy density in Eq. (20) suggesting symmetry about 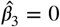 at equilibrium. Thus, we first emphasize that our results are not at equilibrium. Furthermore, the binding terms in the reaction-diffusion equation, Eq. (29), break the symmetry of *δL* with respect to *δL* = 0. This also suggests that the symmetry with respect to 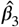 is perturbed. Specifically, the formation of LAT-rich inclusions leads to increased binding in those regions, thus stabilizing them. In contrast, when 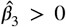 the formation of LAT-depleted inclusions leads to reduced binding in these regions while the density in the LAT-rich phase is also lower than when 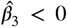, further reducing binding. This lower binding then favors the formation of circular LAT-depleted inclusions.

In Figs. 10a and 10c, initial densities of *L*_0_ = 2.8 and *L*_0_ = 3.2 are considered, respectively. We find that in both cases, there exist parameter combinations at which phase separation is not observed. Phase separation does not occur at small and large values of 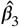 for *L*_0_ = 2.8 and *L*_0_ = 3.2, respectively. These results suggest that the constraint of LAT mass conservation does not make the free energy density minima energetically favorable solutions, thus inhibiting phase separation. For *L*_0_ = 2.8, we now also observe the emergence of circular LAT-rich droplets not seen in the other two phase diagrams, and when *L*_0_ = 3.2, we only observe LAT-depleted inclusions. The latter are non-circular for small values of 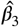 and become circular as 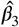 increases. For both Figs. 10a and 10c the arguments for the transition from LAT-rich to LAT-depleted and from circular to non-circular inclusions follow the same reasoning as discussed for Fig. 10b. If we now consider a fixed set of parameter values, 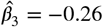 and 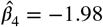, highlighted in red in Fig. 10, we observe that increasing the density changes the morphology from circular LAT-rich inclusions to non-circular LAT-depleted inclusions to circular LAT-depleted inclusions. The same trend can be observed in reconstitution experiments, as shown in Fig. 11 (***Huang et al., 2016***, Fig. 2c).

**Figure 11.**
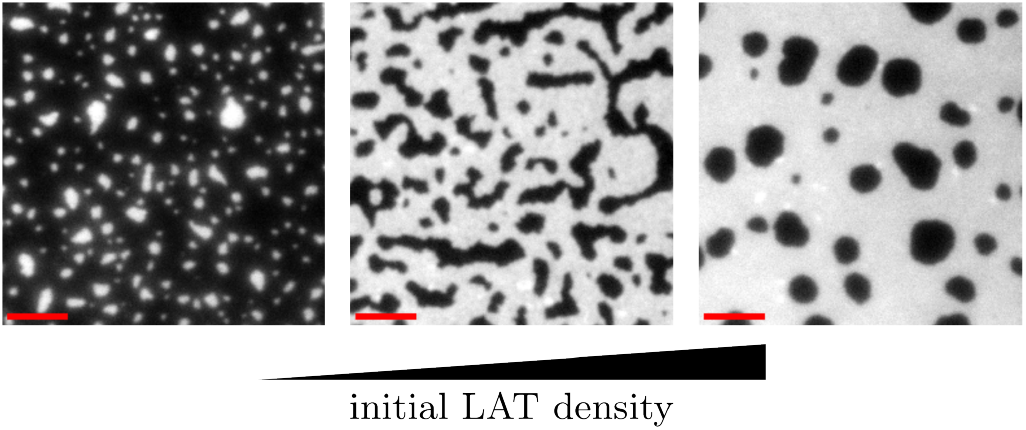
With increasing density, the observed morphologies change from near circular LAT-rich inclusion to elongated LAT-depleted islands to LAT-depleted circular islands. The figure is reproduced from data described by ***Huang et al. (2016***). (scale bars: 5 *μ*m)

**Figure 12.**
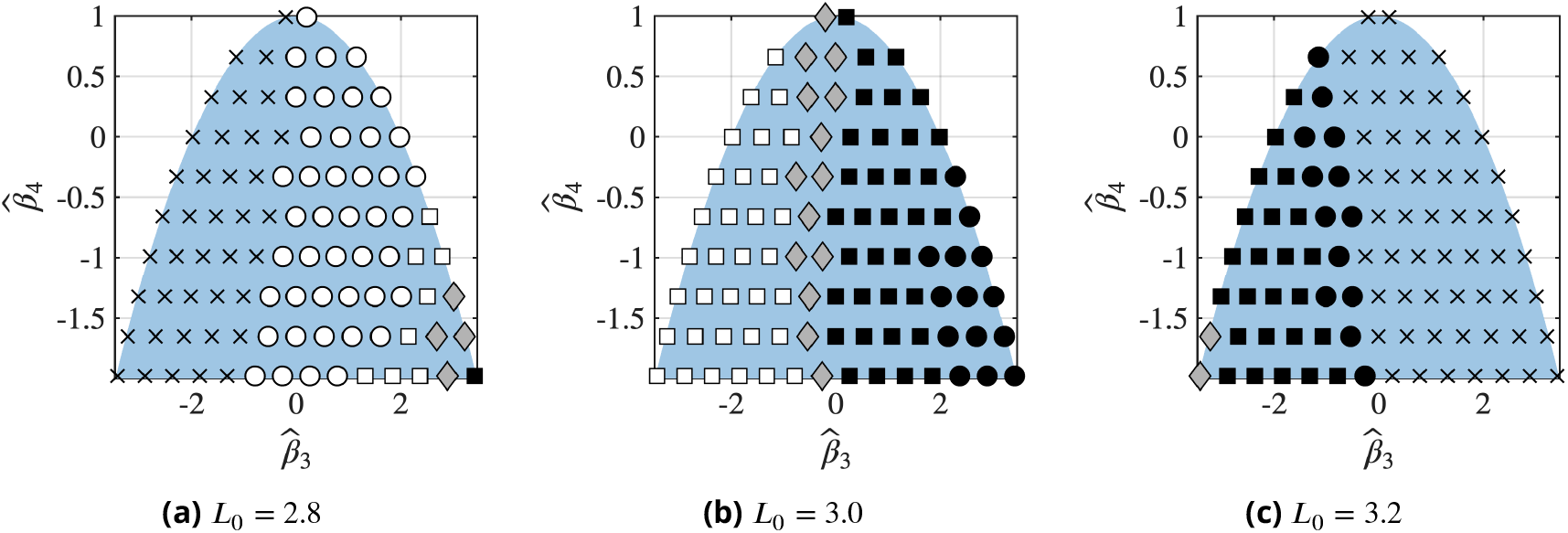
Phase diagrams of the observed morphologies at the end of each simulation for varying free energy parameters 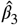 and 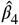, defined in Eqs. (33) and (34), respectively, at different initial densities *L*_0_. The blue background indicates the domain of permissible parameter combinations based on Eqs. (22) and (24). Each marker represents a single simulation and the meaning of each marker is indicated in Fig. 9. The unbinding rate is twice the value of that using in Fig. 10, i.e. 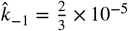.

Next, we investigate how the rate constants in the reaction diffusion equation, Eq. 29, affect the phase diagram in Fig. 10. To this end, we consider an unbinding rate constant of 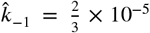, twice the value used for Fig. 10 and plot the corresponding phase diagrams in Fig. 12. These phase diagrams reveal an overall similar structure as those in Fig. 10. Most notably, however, the region where phase separation does not occur expanded in Figs. 12a and 12c compared to Figs. 10a and 10c. This is a result of the lower equilibrium bond density due to an increased dissociation constant, making phase separation less favorable. These observations show that changes in binding rates can affect whether phase separation occurs in experiments even when *ϕ* > *ϕ*_th_. Furthermore, comparison of Figs. 10 and 12 shows that the morphologies are also impacted by the the rate constants: Throughout all phase diagrams, we find small shifts in the location of phase boundaries between different morphologies. This emphasizes that the observed morphologies can also be affected by the binding parameters.

Next, we discuss the physical interpretation of the free energy parameters 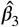 and 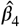, defined in Eqs. (33) and (34). 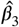 and 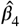 are associated with the cubic and quartic terms of the free energy, respectively, and both multiply the difference *ϕ* − *ϕ*_th_ (cf. Eq. (20)). Thus, they indicate how the free energy landscape changes with changing bond densities. The parameter 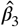 skews the free energy landscape towards either LAT-rich or LAT-depleted minima, indicating that one becomes favorable over the other. This can be understood as cooperative phase separation due to crosslinking—if 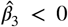 is negative, cooperativity promotes clustering whereas the opposite is the case if 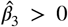. Therefore, we expect 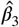 to be affected by, for example, allosteric effects and the LAT valency. In contrast, the parameter 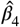 affects how the depth and location of the free energy density minima changes (symmetrically) with changing bond densities. For example, 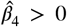 may capture effects such as the inability of a condensate to restructure its bond configuration to yield higher densities, thus getting kinetically arrested. Similarly, if 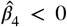, the densities of the energy minima become more dissimilar, indicating the formation of very dense LAT-rich clusters at the expense of a more dilute LAT-depleted phase. Therefore, we expect that 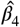 is affected by crosslinker properties such as length, flexibility, bond energy and life time. Furthermore, 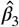 and 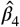 might also be influenced by the lipid composition, which can affect LAT-LAT interactions (***Chung et al., 2021***; ***Wang et al., 2023***). Finally, we note that the above discussion is purely qualitative and that the inhomogeneous LAT and bond densities as well as the constraint of mass conservation render a mapping of the phenomenological free energy parameters to measurable physical parameters of the system challenging. Thus, a deeper understanding of such mappings requires further experimental and theoretical studies to enable predictive modeling of condensate behavior from molecular details.

## Conclusion

In this article, we presented two models that address four open questions introduced in the Introduction section regarding reconstitution experiments of LAT condensates on supported lipid bilayers. (1) Using the Smoluchowski aggregation model, we focused on the early times of the experiments and showed that a large number of small LAT clusters is required before the rapid formation of larger clusters can occur. The formation of small cluster leads to an onset time that is regulated by the binding probability between individual monomers or small oligomers. This binding probability is expected to be low since bonds form with the help of three crosslinking molecules binding from the bulk fluid, leading to potentially long lag times. Furthermore, the binding probability might be affected by other system parameters, including LAT valency and the LAT and crosslinker concentrations. We then used this insight to propose a field-theoretic model to gain a deeper understanding of the spatio-temporal dynamics of phase separation of LAT. This model further confirmed the dependence of the onset time on the LAT valency and effective crosslinker binding rates. (2) Using the field-theoretic model, we showed that the distinct experimentally-observed morphologies arise from the same underlying dynamics, modulated by variations in the free energy parameters, binding rates and initial LAT concentration. This finding provides a unifying explanation for the diversity of morphologies seen across experimental realizations. (3) Furthermore, we found that the rapid emergence of near-stationary patterns upon phase separation can be explained by the reduced diffusivity within the LAT-rich phase. (4) Finally, our model showed that phase separation is suppressed in some parameter regimes. Importantly, this is not only observed for high dissociation constants but also various other combinations of free energy parameters and binding rates. This result explains why experimental realizations may fail to show phase separation altogether under some experimental conditions.

While the models presented in this article describe the phase separation behavior of LAT in reconstitution experiments, they are sufficiently general to also apply to the interaction of other multivalent proteins on membranes. For example, the epidermal growth factor receptor (EGFR), a receptor tyrosine kinase, forms protein condensates through interactions with Grb2:Grb2 or Grb2:SOS:Grb2 as crosslinkers. These interactions are mediated by the phosphorylation of tyrosine residues in the EGFR cytoplasmic tail (***Lin et al., 2022b***; ***Phan et al., 2024***). Like LAT, this condensation process exhibits a potentially large lag time (***Phan et al., 2024***) and shows the same trends in the dependence of the observed patterns on the EGFR density (***Lin et al., 2022b***, Fig. 1c). A similar example is fibroblast growth factor receptor 2 (FGFR2), which phase separates in the presence of its downstream effectors, SHP2 and PLC*γ*1, in response to receptor phosphorylation. However, the morphologies, patterns and time dependence have been investigated in less detail than for LAT and EGFR (***Lin et al., 2022a***). Another example is the adhesion-receptor nephrin, which can cluster on membranes through interactions with Nck and N-WASP, driven by nephrin phosphorylation. Similar to LAT, the observed patterns and morphologies are concentration dependent and the final structures are near stationary (***Banjade and Rosen, 2014***). The examples of EGFR, FGFR2, and nephrin thus suggest that the results of this article are universal and could generally apply to phase separation of multivalent proteins on lipid membranes.

## Acknowledgments

We thank L.J. Nugent Lew for his assistance in gathering, interpreting, and postprocessing the experimental data used in this study. We also gratefully acknowledge William Y. C. Huang for his previously published work pertaining to this article. Research reported in this article was supported by National Institute of Allergy and Infectious Diseases of the National Institutes of Health under award number 2P01AI091580-11A1 and by the Novo Nordisk Foundation under the Center for Geometrically Engineered Cellular Systems (NNF17OC0028176).

## Footnotes

1 The terms *lag time* and *delay time* are not to be confused with the delay time observed in live cells between TCR-pMHC binding and LAT condensation. There, the delay originates from both phosphorylation of LAT and subsequent crosslinking.

2 The Brownian kernel in Eq. (7) is often referenced in the more general form *k*_*ij*_ ∝ (*R*_*i*_ + *R*_*j*_)^*d*−2 (^*D*_*i*_ + *D*_*j*_), where *d* denotes the spatial dimension and *R*_*i*_ is a measure of the radius of clusters of size *i* (***Jullien, 1992***; ***Odriozola et al., 2001***; ***Moncho-Jordá et al., 2004***). However, the original derivation makes the provision *d* > 2 (***Meakin, 1992***; ***Krapivsky et al., 2010***). Yet, dimensional analysis and an exit time analysis indicates that the characteristic time indeed takes the form of the rate constant in Eq. (7) (***Krapivsky et al., 2010***).

3 It is well-understood that the fractal dimension depends on whether aggregation is diffusion- or reaction-limited (***Meakin, 1992***). Thus, the effective fractal dimension might vary with time or might be ill-defined in our simulations, suggesting that the exponent *γ*_H_ could vary over the course of our simulations. However, we show in the SM (Fig. S1) that our results are not sensitive to the choice of *γ*_H_.

